# Isolation and characterizations of a novel H6N1 avian influenza virus from Von Schrenck’s Bittern (Ixobrychus eurhythmus) in Jiangxi province, China

**DOI:** 10.1101/2021.06.24.449698

**Authors:** Hamidreza Attaran, Wen Jin, Jing Luo, Chengmin Wang, Hongxuan He

**Author notes:** **Corresponding author: Hongxuan He and Hamidreza Attaran:** National Research Center for Wildlife-borne Diseases Institute of Zoology, Chinese Academy of Sciences, Beijing, China.

## Abstract

In June 2013, the first case of human infection with an avian H6N1 virus was reported in a Taiwanese woman. Although this was a single non-fatal case, the virus continues to circulate in Taiwanese poultry. As with any emerging avian virus that infects humans, there is concern that acquisition of human-type receptor specificity could enable transmission in the human population. Despite mutations in the receptor-binding pocket of the human H6N1 isolate, it has retained avian-type (NeuAca2-3Gal) receptor specificity.

An H6N1 AIV was isolated from a Von Schrenck’s Bittern during national active surveillance project for avian influenza viruses (AIVs) in wild birds in Jiangxi province, China 2018. Phylogenetic analysis showed that this strain received its genes from H1, H2, H3, H4, H6 and H10 AIVs in different places. This strain was found to be minimally pathogenic in mice and was able to replicate in mice without prior adaptation. Considering that the reassorted H6N1 virus was isolated from Von Schrenck’s Bittern in this study, it is possible that this bird can play an important role in the generation of novel reassorted H6 AIVs.

In this study, H6N1 virus is a wild virus migration brought from different regions of the AIV gene into the natural gene pool, resulting in the production of a recombinant new virus. The H6N1 virus can be seen as a link in the evolution of the virus and the evolution of other viruses.

## Introduction

Influenza A viruses belong to the Orthomyxoviridae family. An influenza A virus consists of an enveloped, negative-strand RNA with eight gene segments, which can adapt to human stably, and lead to sustained transmission between humans[1, 2] causing mild or severe diseases[3].

Based on the antigenic properties of the hemagglutinin (HA) and neuraminidase (NA) glycoproteins, influenza A viruses are classified into 18 HA and 11 NA subtypes [4, 5].

Most subtypes of avian influenza viruses (AIVs) have been identified in aquatic birds, which are considered a natural reservoir for AIVs[6]. Though aquatic birds, including domestic ducks, do not usually display clinical signs. When aquatic birds are infected with AIVs, they provide a niche for the reassortment of low pathogenic avian influenza viruses. They can serve as progenitors of highly pathogenic avian influenza viruses [7, 8].

Previous pandemics in humans were mainly caused by birds and required a shift from α2, 3-linked sialic acid receptors (avian receptor) to α2-6-linked sialic acid receptors (human type receptor) [9]. Therefore, understanding the potential of new AIVs that obtain human receptor specificity is a key factor in assessing potential pandemic threats. For a long time efforts to monitor and control avian influenza all over the world have focused on the H5, H7 and H9 subtypes, because of the high mortality rate and major economic losses in affected flocks. However, it is difficult to predict the subtype of AIV that can cross the species barrier and infect humans in the future. More and more evidence suggests that low pathogenic AIVs (LPAI) are also important, mainly because they could infect poultry and humans if they undergo reassortment to produce pathogenic forms [10, 11].This possibility was manifested in 2013 by the unexpected human cases of low pathogenic avian influenza H7N9 viruses in China; H7N9 has resulted in more than 700 deaths in humans ∼ 36% [12].

The first H6 influenza virus was isolated from Turkey in 1965 and since then the H6 virus has been increasingly isolated from wild, domestic aquatic, and terrestrial bird species throughout the world.

The H6 subtype avian influenza viruses were more promiscuous in host range than other AIVs subtypes [13]. The H6 subtype is the most common subtype in domestic ducks in Southern China [14, 15].

Previous studies have shown that H6 AIVs were involved in the generation of human H5N1, H9N2 and H5N6 viruses [16, 17]. The H6 reassortant viruses can cross the species barrier thus infecting mammals including humans without adaptation. For example, H6N6 AIVs were isolated from swine in Southern and Eastern China with clinical symptoms [18, 19]. A H6N1 virus was also isolated from a dog in Taiwan [20]. Recent seroprevalence studies have shown that the H6 viruses were seropositive in 19 provinces of China among occupational exposure workers.[21] The H6 specific antibody was significantly elevated in veterinarians in the United States [22]. In May 2013 an H6N1 virus was isolated in Taiwan from a 20 year old woman whose symptoms included cough, fever, muscle ache, and headache [23].

The study of H6 viruses isolated from Taiwan in the past 14 years suggests an elevated threat of H6 viruses to human health [24].

These data suggest that the H6 subtype avian influenza virus may pose a potential public health threat. In this study, we investigated influenza H6N1 subtype viruses isolated in mainland China for genetic and antigenic characteristics, receptor-binding properties and kinetics of infectivity in mammalian.

## Materials and Methods

### Ethics statement

The study was carried out in strict accordance with the Guide for the Care and Use of Laboratory Animals of the Ministry of Science and Technology of the People’s Republic of China. The protocol was approved by the Committee on the Ethics of Animal Experiments of Institute of Zoology, Chinese Academy of Sciences (CAS). (Approval number IOZ20160046 for mice).

### Sampling

The environmental samples were collected from Jiangxi province. All the sampling sites are located in East Asian–Australian migratory flyway, from Southeastern Siberia to Eastern Asia and Australia. The sampling sites consisted of important stopovers, breeding locations and wintering for migratory birds. The selected sampling sites were areas at which the local staff in bird-banding and wild bird rescue was known to the local population and thus had access to the wild birds.

### Virus isolation and sequencing

After filtering the specimens they were inoculated into a 10-day-old specific-pathogen-free embryonated eggs allantoic cavity (Beijing Merial Ltd). After incubation for 72□hours at 37□°C, the allantoic fluid was collected and tested using hemagglutinin (HA) and hemagglutinin inhibition (HI) assays. In order to confirm positive samples reverse transcription–PCR was done.

### Sequencing and phylogenetic analysis

All eight gene segments of the isolated H6N1 virus were amplified (NEB), purified (Omega) and sequenced (BGI, China), then compared with reference viruses in Genbank (http://www.ncbi.nih.gov) and GISAID. Phylogenic trees with molecular evolutionary genetics analysis (MEGA 6) were constructed for each gene segment.

### Animal study

To evaluate pathogenicity of the H6N1 isolated virus in mammalian hosts, 26 female BalB/C mice were inoculated intranasally with 50 μl H6N1 isolate (1×104 EID50) and PBS respectively. Three mice were euthanized on the 3rd AND 5th day post inoculation respectively. Their brain, liver and lungs were separated for TCID50 assay and RNA extraction used in real time qPCR. The remaining mice was observed over a 14 day period following inoculation for weight losses and their survival rate.

### Histopathologic analysis

The lung tissues of euthanized mice were fixed in a 10% phosphate-buffered formalin, embedded in paraffin, cut into 4-mm and stained with hematoxylin and eosin (H&E) for histopathological changes.

## Results

### Homology analysis of H6N1

The H6N1 gene sequencing results using the Snapgene software revealed that the HA gene was 1709bp in length and encoded 567 amino acids. The length of the NA gene was 1433bp and encoded 469 amino acids. The length of the PB2 gene was 2280bp, and encoded 759 amino acids. The full length of the PB1 gene was 2309bp and encoded 757 amino acids. The PA gene was 2233bp in length and encoded 716 amino acids. The full length of the NP gene was 1550bp, and encoded 498 amino acids. The total length of the M gene was 1031bp and encoded 252 amino acids. The full length NS gene was 985bp and encoded 230 amino acids.

The NCBI Blast (http://blast.ncbi.nlm.nih.gov/Blast.cgi) analysis revealed that the isolates of H6N1 was isolated from countries and regions of different migratory routes. These routes included Eastern Asia and Western Europe and were reorganized by different subtypes (see Table 1 (A)).

**Table 1.**
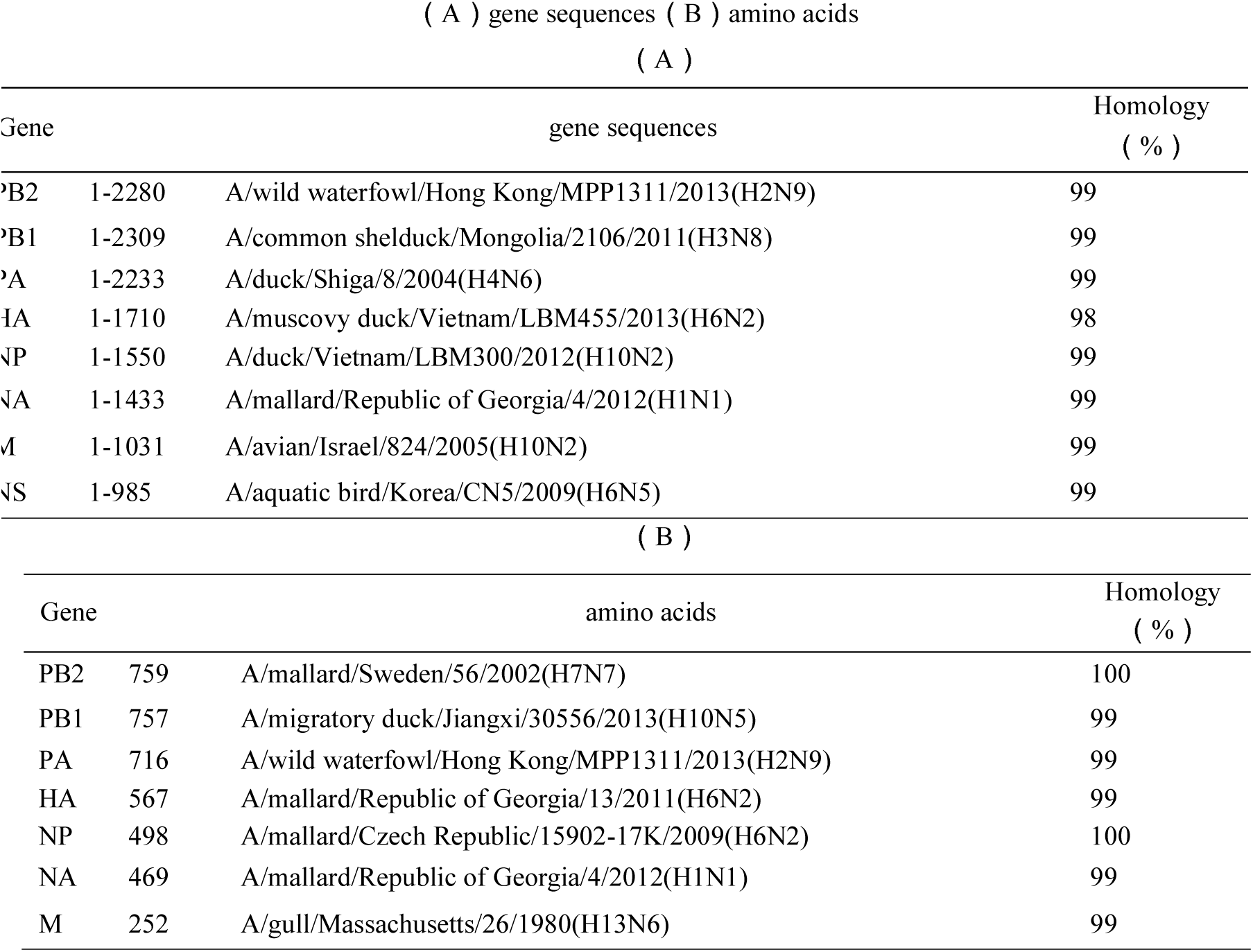
Homology analysis of gene sequences and proteins of H6N1

The highest level of similarity was at 98% between the HA gene and the Vietnamese isolate A/Muscovy duck/Vietnam/LBM455/2013 (H6N2). The NP nucleic acid fragments was 99% similar the H10N2 isolates from the Vietnam live poultry market in 2012. The PB2 gene was 99% similar to the PB2 gene of H2N9 strain isolated in Hong Kong, China in 2013.

The HA, Np and PB2 gene fragments were found in the South of China and Southeast Asia. The most similar sequence of these gene fragments were found in Vietnam and Hong Kong. Vietnam and Hong Kong are a part of China’s eastern migratory flyway. This flyway is part of the East Asian - Australian migration route.

The PB1 gene was 99% similar to the H3N8 strain found in 2011 of a wild duck in Mongolia. The PA gene was 99% similar to the H4N6 isolated genes of ducks from Shiga, Japan in 2004. The NS gene was 99% similar to the 2009 NS fragment of the Korean H6N5 strain. Mongolia, Japan and South Korea are part of the East Asian-Australian Bird Migration route. The NA gene was 99% similar to the nucleotide sequence of the H1N1 virus isolated from the Mallard in 2012 located in the Republic of Georgia, Eastern Europe. The M gene was 99% similar to Avian H10N2 virus isolated in 2005 from Israel in the Middle East. The Republic of Georgia and Israel are part of the West Asia-East African migratory flyway.

The amino acid sequence of the H6N1 isolate was analyzed by using the NCBI Blast application. As shown in Table 1 (B), the amino acid of the viral protein was similar to different subtypes. Thus proving the researched isolated H6N1 virus is a novel recombinant virus.

## Evolutionary analysis of H6N1

### The HA Gene

The HA gene can be divided into Group I, Group II and Group III large lineage [25]. This sorting is based on evolutionary analysis of wild and domestic H6 subtype viruses in China, as shown in Figure 1.

**Figure 1:**
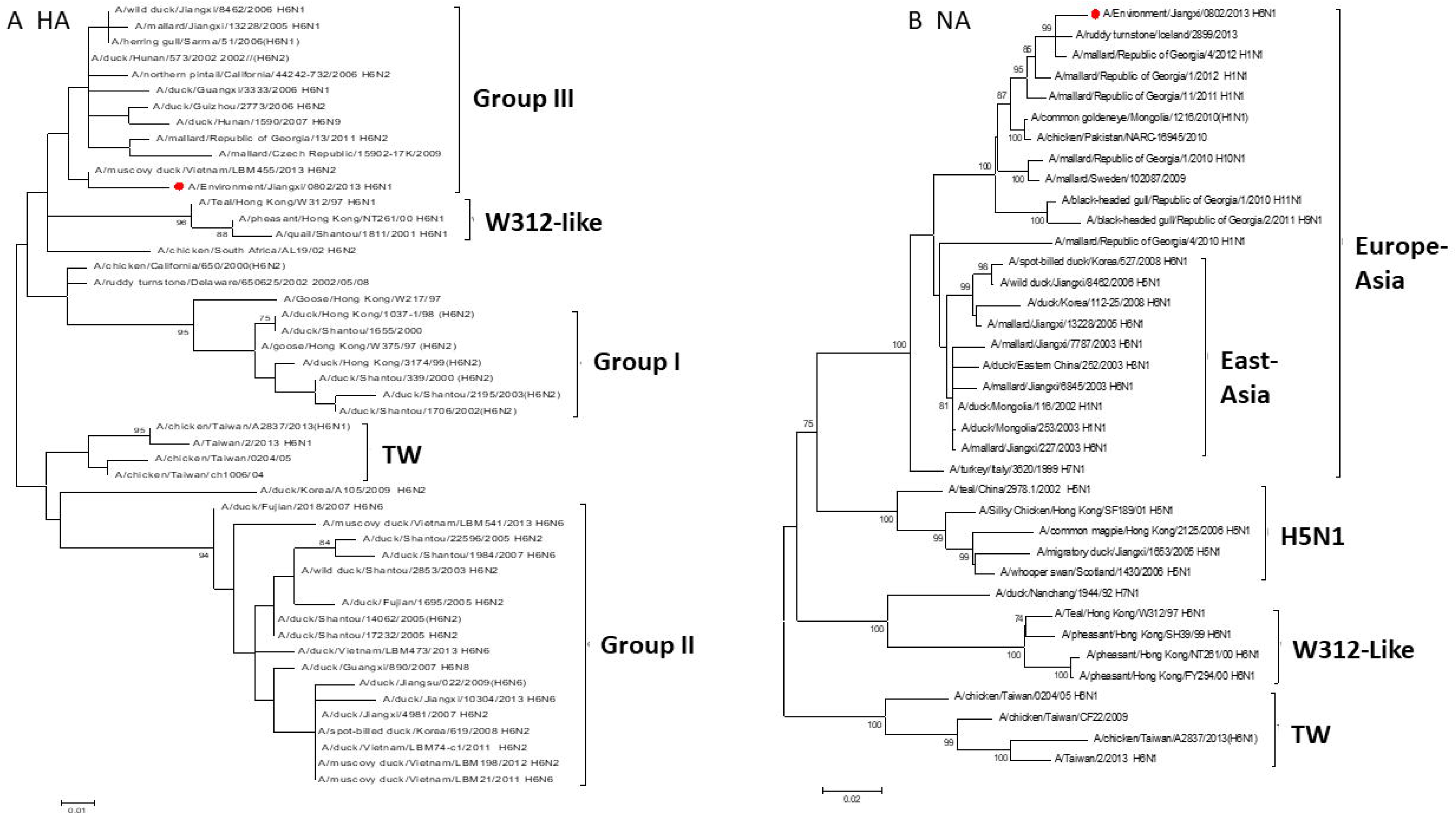
Demonstrates the phylogenetic relation of Hemagglutinin (A) and neuraminidase (B) of the H6N1 virus. The phylogenic tree represents the H6N1 virus analyzed in this study. The phylogenic tree was constructed using the Maximum Likehood method of MEGA 5.0. The reliability of the phylogenic trees was assessed by bootstrap analysis with 1000 replications. The following represented phylogenic trees were also constructed using the same methods.

The Group I lineage consisting of DK/HK/1037-1/98 and A/Duck/Shantou/339/2000 H6N2 are representative of one lineage, mainly from Guangdong, Shantou and Hong Kong isolates. The Group II lineage consisting of A / Duck / Shantou / 2853/2003 H6N2 are mainly composed of viruses isolated from Guangdong and Fujian. The Group II virus strain was popular in domesticated ducks from 2002 to 2005. The Group III lineage consisting of A / Duck / Hunan / 573/2002 H6N2 are mainly composed of Jiangxi and Hunan isolates. The Group III virus strain is similar to wild birds or migratory birds, as well as poultry viruses in other countries. This virus has a high probability of recombination between wild birds and poultry, belonging to the European gene pool [25].The H6N1 strain isolated in this study is located in the Group III lineage. Our H6N1’s evolutionary pattern was similar to the 2013 H6N2 isolate in Vietnam and the 1997 Hong Kong H5N1 patient isolates. The A/Hong (HK/97) Kong/156/97 H5N1, A/Hong A/Teal/Hong Kong/W312/97 provides a gene fragment for the H6N1 virus [16] (W312-like), however it does not belong to the same lineage.

### The NA Gene

The evolutionary analysis of the NA gene is shown in Figure 1. The H6N1 virus isolated from minor poultry in southern China is similar to the W312-like cluster virus However the NA gene of the isolated H6N1virus in this experiment is not present in the W312-like cluster virus. Our H6N1 virus and H6N1 human isolated virus A / chicken / Taiwan / A2837 / 2013 (H6N1) (A2837) both belong to the Eurasian branch. The NA gene may be brought into China by the migration of wild birds. The NA gene’s code is most similar to the 2012 H1N1 virus isolated in the Republic of Georgia.

### The PB2\PB1\PA\NP\M\M\NS Genes

Phylogenetic analysis of the PB2 gene revealed that the PB2 gene of the H6N1 strain was located in the Group III lineage. The Duck/Hunan/573/02 virus cluster in H6N2 and the HA cluster in Group III both belong to the European gene library.

Most of the wild or domestic animal isolates from the southwestern provinces of China, Jiangxi, Hunan and Guangxi are part of the Group III gene. In addition the PB1, PA and PB2 genes are located within the Group III European gene library. Figure 2 (A) - (C) demonstrates the distant evolutionary relationship of the southern

**Figure 2:**
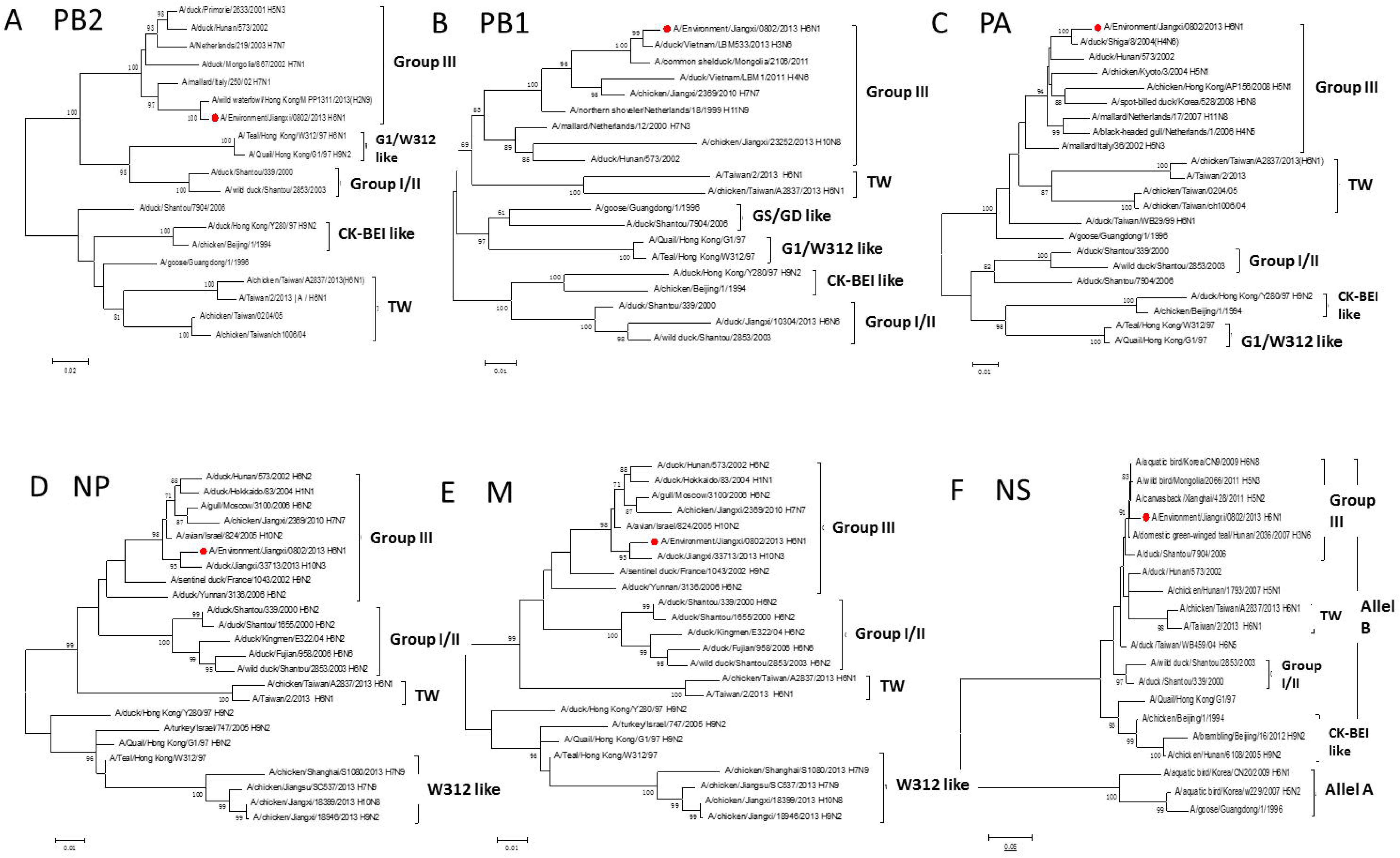
The phylogenic tree shows the relationship between the inner protein genes of the H6N1 virus. (A)PB2, (B)PB1, (C)PA,(D)NP,(E)M,(F)NS genes

W312-like (CK/Bei-like), A/chicken/Beijing/1/1994H9N2, A/goose/Guangdong/1/1996 H5N1Gs/GD-like) and A/chicken/Taiwan/A2837/2013 strain H6N1 (A2837/TW). The results show that the RNA polymerase PB2, PB1 and PA genes of H6N1 isolates of wild birds and domestic birds evolved from the European branch gene pool.

Figure 2 (D) - (F) shows the reference strains evolutionary relationship between M, NS and NP gene. Furthermore Figure 2 (D) - (F) demonstrates the distant evolutionary relationship of the M, NS and NP gene in relation to the W312-like, CK / Bei-like, G1-like, Gs / GD-like and A2837 viruses. Thus in conclusion 8 gene segments of H6NI isolates from the Eurasia AIV gene pool evolved.

## H6N1 key AA site analysis

### H6N1 HA and NA protein analysis

The ORF region was searched using the Snapgene software. The amino acid sequence was also deduced using the Snapgene software. (SEQ ID NO: 324-330) The amino acid sequence of the H6N1 isolate was PQIETR ↓ G, and there was no continuous basic amino acid insertion. Thus we discovered the H6N1 isolate was a low pathogenic strain. We evaluated the receptor site sequence analysis of the A138, E190, L194, G225, Q226 and G228 and Glutamine 226. This analysis revealed that the H6N1 virus can easily bind to the alpha -2, 3-SA receptor. Studies have shown that S138A, I155T, T160A and H183N mutations of HA protein can increase the affinity of virus to alpha -2, 6 receptor [26, 27]. As shown in table 2(A), the 138 and 160 amino acids of the HA protein of H6N1 are Alanine. This causes the virus to contain the possibility of binding to the mammalian receptor, posing a potential risk to the mammal. The analysis of the NA protein sequence was conducted using the Snapgene software. The amino acid deletion in the stem ring of NA protein, increased the pathogenicity of the virus in mice [28]. Furthermore our analysis revealed there was no amino acid deletion in the ring structure of the NA protein of the isolate H6N1. The amino acid 275 of NA protein is a neuraminidase inhibitor drug-sensitive site. The H275Y makes the virus drug resistant [29]. The amino acid of the H6N1 isolate at the 275 position is Histidine. Thus indicating that the strain is sensitive to Oseltamivir and is not drug resistant.

**Table 2(A):**
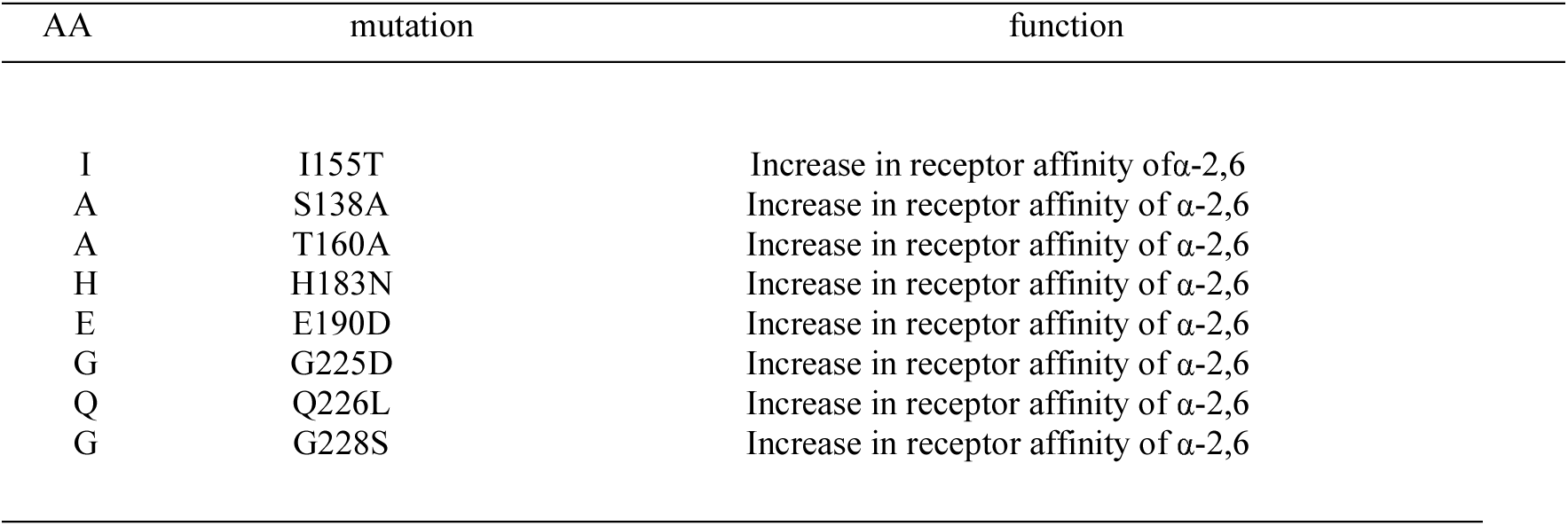
Selected characteristics of Amino Acids of hemagglutinin in the H6N1 virus

**Table 2(B):**
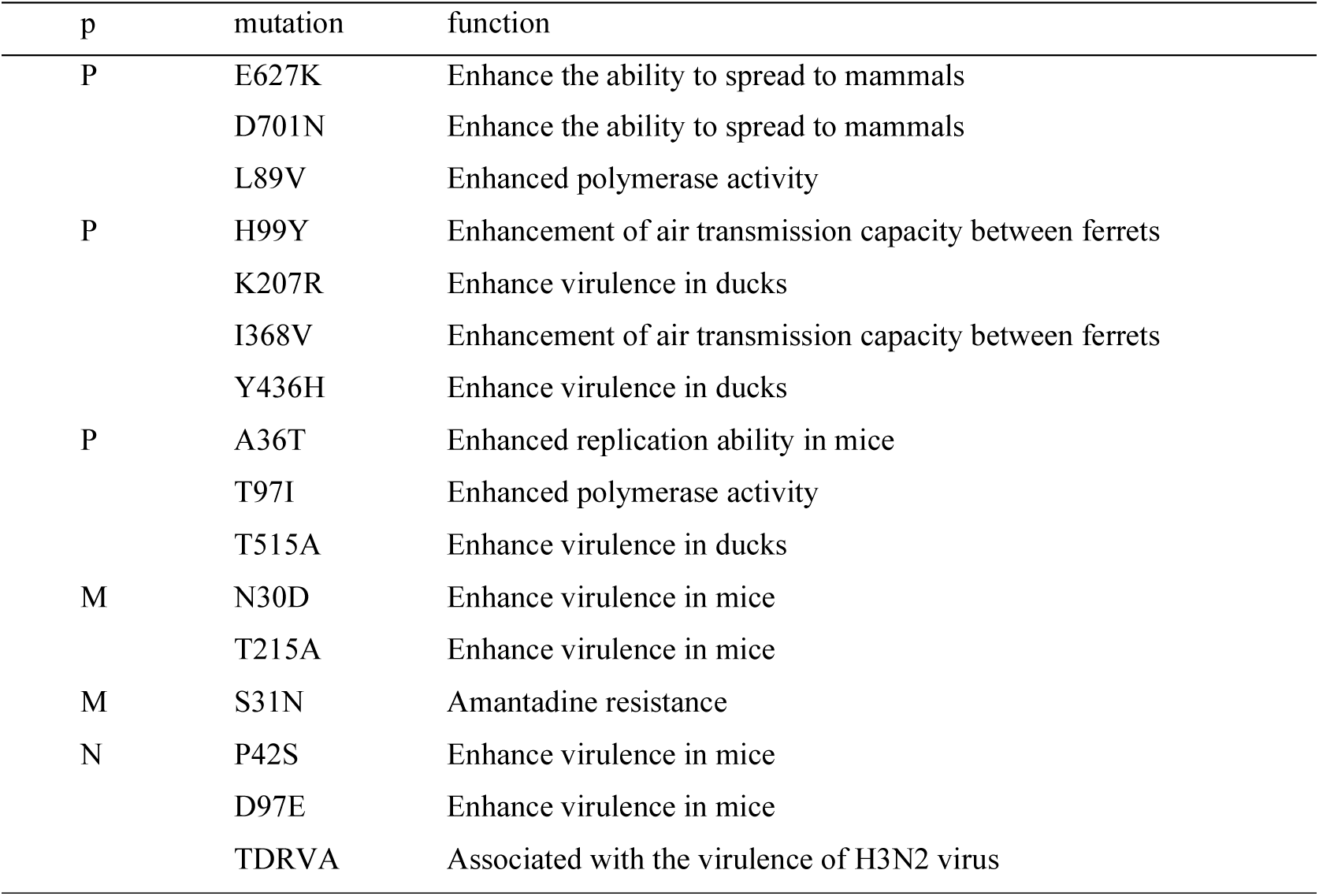
Selected characteristic amino acids of inner proteins of the H6N1 virus

### HA, NA protein glycosylation site analysis

The HA and NA protein potential glycosylation sites of the H6N1 virus were analyzed using the glycosylation online search tool. (HANetNGlyc 1.0 Server http://www.cbs.dtu.dk/services/NetNGlyc/) The HA protein has five glycosylation sites. The results are shown in Table 3(A). The HA1 protein had four glycosylation sites, 27 (NSTT), 39 (NVTV), 182 (NNTG) and 306 (NKTF). The HA2 protein contained only the 498 (NGTY) glycosylation site. In addition the glycosylation sites of the NA protein are shown in Table 3(B). The table includes the 50 (NQSI), 58 (NNTW), 63 (NQTY), 68 (NISN), 88 (NSSL), 146 (NGTV) and 235 (NGSC) NA glycosylation sites

**Table 3 (A):**
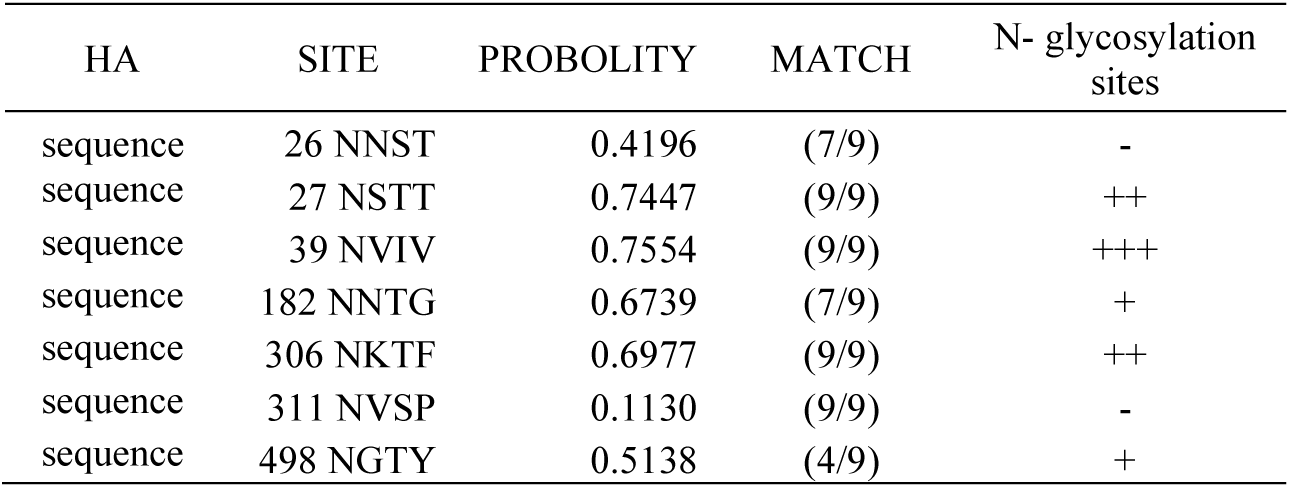
Comparison of the potential glycosylation sites in the amino acid sequence of the HA protein

**Table 3 (B):**
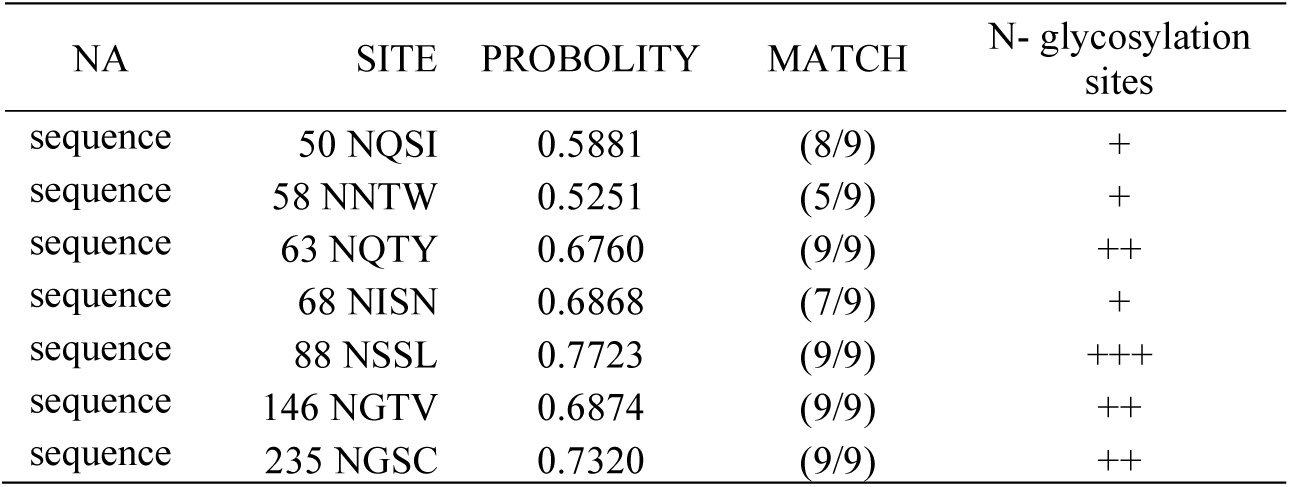
Comparison of the potential glycosylation sites in the amino acid sequence of NA protein

### H6N1 internal protein analysis

The key amino acid mutations in internal proteins also lead to changes in viral replication ability and virulence, as shown in table 2(B). The mutation of the Glutamate to a Lysine (E627K) combined with the mutation of the Aspartic Acid to the Asparagine (D701N) plays an important role in the transmission of H5N1 AIV to mammals. [30-32] these two mutations are often found in H7N9 isolates [33, 34].

The 627 position of the H6N1 isolate was an E amino acid. The 701 position of the H6N1 isolate was a D amino acid. Thus revealing the minimal infection ability of the H6N1 isolate on mammals.

The mutation of the 89 lysine to valine in the PB2 protein (L89V), enhanced the viral polymerase activity [27]. In this study the H6N1 strain was valine in the 89 position, which can enhance the polymerase activity of the strain. The H99Y, R207K and H436Y mutations in the PB1 protein increased the virulence of the H6N1 virus in ducks and the transmission ability among ferrets [35, 36]. The T515A mutation of the PA protein can enhance the toxicity of the H6N1 virus in ducks. Furthermore the T97I mutation can enhance the activity of the virus’s polymerase [37]. In addition the A36T mutation can enhance the virulence of the H6N1 virus to mice. [35, 36]

The PB1 and PA proteins of the H6N1 isolate showed no mutation at the listed positions. The Polymerase Chain Reaction (PCR) analysis of H6NI’s PB1, PB2 and PA proteins revealed only the L89V mutation in the polymerase. The L89V mutation located in the PB2 protein does not infect mammals. Amantadine and Rimantadine are anti-influenza drugs designed for the M Gene Ion Channel. An Amantadine drug resistance can be formed when the serine mutates to an aspartic acid (S31N). [38]

The isolated H6N1 M2 protein located in the 31 position is serine, revealing no form of drug resistance. However mutations in the M1 protein’s N30D or T215A positions can increase the virulence of the virus in mice [35]. For the NS1 protein, the amino acid mutation sequences in D42S, D97E, and TDRVA127N can cause viral virulence enhancement [35].The NS protein of the H6N1 isolate was serine in position 42, was a glutamic acid in position 97, and was an asparagine in position 127. These three positions were mutated, thus enhancing the toxicity of H6N1 in mice. In conclusion the mutated amino acids of the H6N1 virus in the M1 and NS1 proteins can enhance the virulence of the virus.

### H6N1 α-2, 3-SA analysis

The viral receptor specificity plays an important role in the host restriction of an influenza virus. The structure of receptors in upper respiratory tract proteins of mammals and birds is different. AIV mainly recognizes α -2, 3-SA receptors. Whereas the human influenza viruses mainly recognize α -2, 6-SA receptors. [39]

Figure 3 shows H6N1 surface protein receptors using solid-phase binding α -2, 3 or α -2, 6 sialic acid oligosaccharide chain. However the H6N1 isolate specifically binds to α -2, 3 receptors and does not bind to α -2,6 receptors. This explains why the strain has a weak ability to infect mammals.

**Figure 3:**
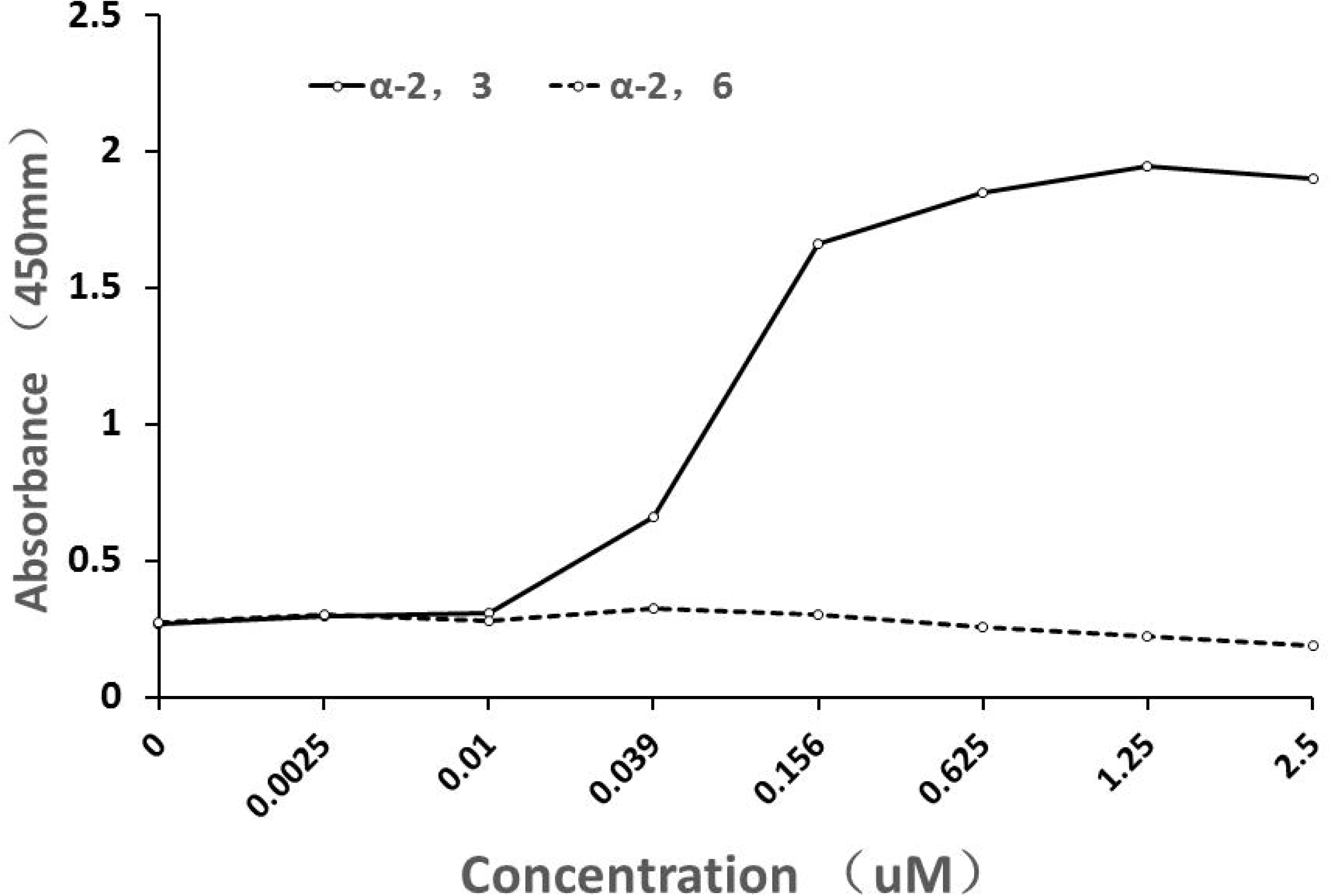
The receptor binding property of hemagglutinin to the H6N1 virus

### H6N1 infection in mice

#### H6N1EID50

By measuring the Egg median infective dose (EID50) of the chicken embryo virus the virus viability was determined. The log10EID50/100uL of the H6N1 isolate was 4.63.

### H6N1 Clinical symptoms in mice

The mice were intranasally inoculated with a 50uL 4.63 EID50 sterile H6N1 virus. As seen in Figure 4 (A) the following day the mice began to lose weight, trembled, clustered together, showed decreased activity, poor response, curled up and got vertical hair. As the days progressed the symptoms increased reaching the highest level (81.6%) on the fourth day.

**Figure 4:**
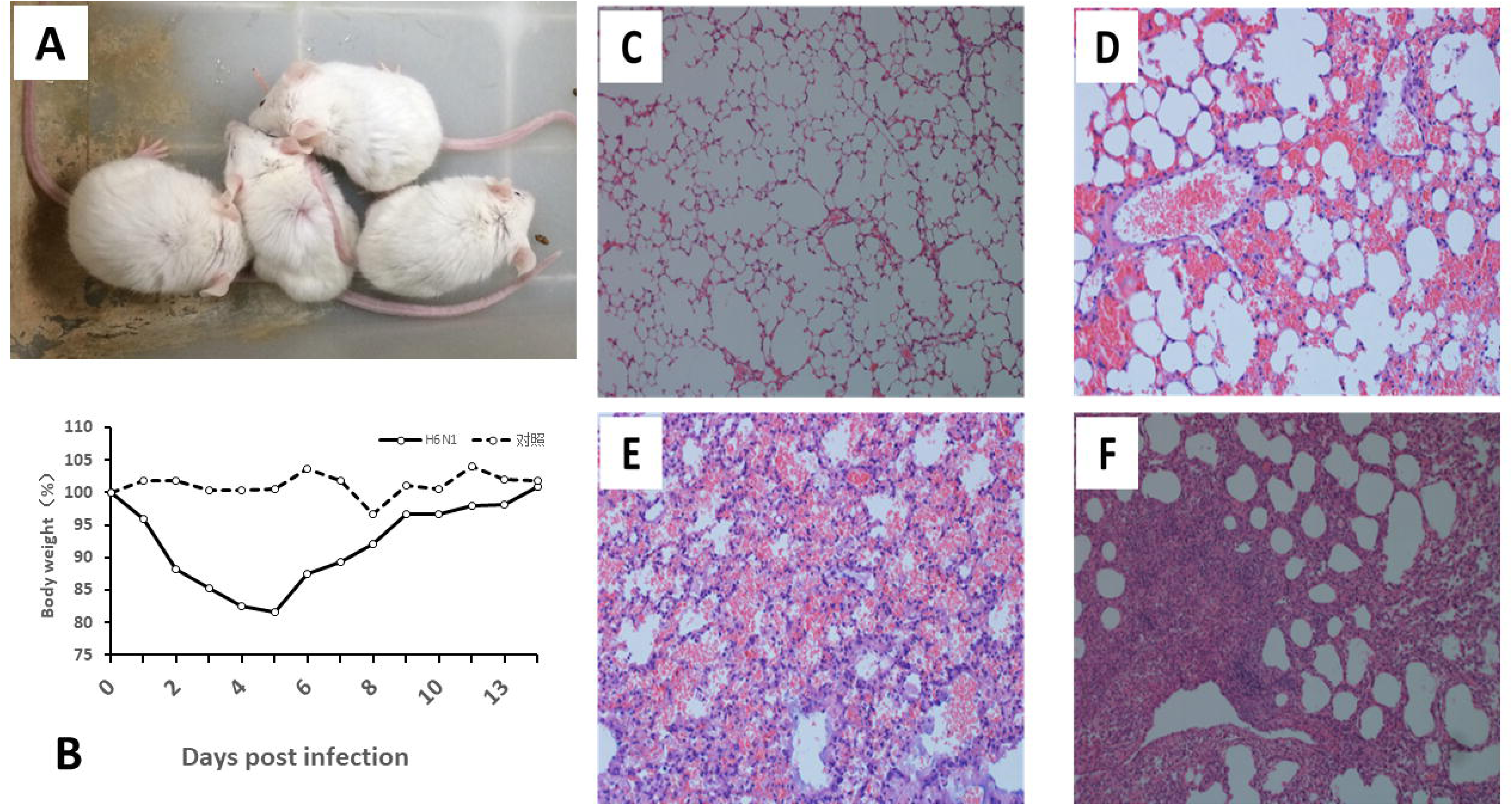
Clinical symptoms in mice inoculated with the H6N1 virus (A) The infected mice showed clinical symptoms, such as curled up, stay together and so on.(B) Body weight changes in two weeks after incubated with H6N1(n=5). Pathological changes in lung: (C) Control(100×) (D) Pneumorrhagia in 3 days post infenction (E) Pulmonary hemorrhage and inflammatory cell infiltration in 5 days post-infenction (F) The Inflammatory cells disappeared and the lung mesenchymal cell proliferated in 14 days post-infenction(200×)

**Figure 5:**
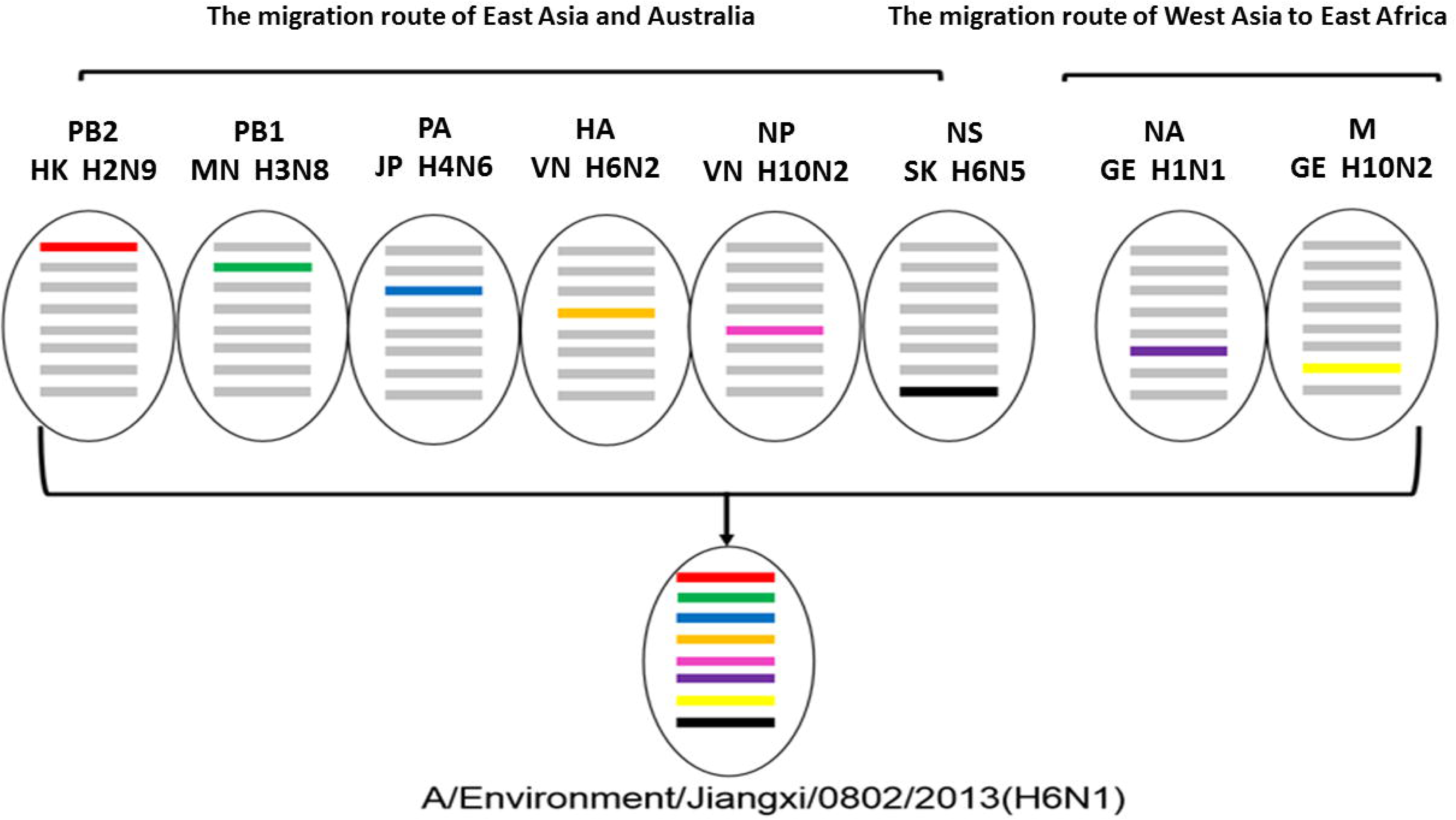
Recombination of genes of H6N1

The mice showed signs of improvement on the fifth day, such as a gradual increase of weight. On the seventh day their level of activity began to improve. On the ninth day the mice returned to their normal level of activity. Furthermore they did not cluster together anymore and their hair returned to its normal state. By the fourteenth day the mice’s clinical signs were stable. However the autopsy revealed the mice’s lungs had partial hemorrhagic lesions and edema. This proved that the inflammation was still not fully cured.

As shown in Figure 4 (B) the mice’s body weight changes were related to their clinical symptoms. The weight of the mice inoculated with PBS in the control group was stable.

### H6N1 In vivo replication

After the inoculation 3 mice were randomly selected and euthanized on both the 3^rd^ and 5^th^ day. Then the mice’s, brain, liver, spleen, lungs and kidney were collected for testing for the H6NI virus in their organs. The organs were tested using the fluorescence quantitative PCR method, the mouse beta –actin and NP genes as internal influenza virus. Influenza viruses were detected on the 3^rd^ and 5^th^ day lung samples.

### However no viruses were detected in their other organs

The virus level of the mice’s lungs was higher on the 5^th^ day compared to the 3^rd^ day. The titer of influenza virus in the lungs was determined using the TCID50 method.

The TCID50 titer of influenza virus log10 in the mice’s lungs was 3.86 + 0.68 on the 3^rd^ day and 6.60 + 0.68 on the 5^th^ day. This result is consistent with the fluorescence quantitative method result. The real-time quantitative PCR experiments and TCID50 titers results reveal that the H6N1 virus can effectively replicate and cause pneumonia in the lungs of mice. The titer of the H6N1 virus on the 5^th^ day was higher compared to the 3^rd^. Which also corresponded to the changes of clinical symptoms and body weight in mice.

### H6N1 the pathological changes of lung

Figure 4 represents the pathological changes of the hematoxylin-and-eosin-stained lung tissues in mice after being inoculated with the H6N1 viruses.

The genomic composition of the isolated H6N1 virus with possible donor viruses. The NA and M genes were possible from the places in the West Aisa-East Africa flyway and the other genes were from East Aisa-Australia flyways.

## Discussion

In 1965 the H6 subtype was first identified in a turkey located in Massachusetts. In the live poultry market of Southern China the first signs of the H6 virus subtype were identified in 1997 [40]. In Southern China during the years of 1999 to 2005 the W312-like H6N1 was prevalent and reorganized in minor poultry [25]. Furthermore in 2000 the A/teal/Hong Kong/W312/97 (H6N1) may have provided a gene fragment for the infection of the avian virus A / HongKong / 156/97 (H5N1) (H5N1) strain in humans [41]. It was confirmed that the H6 virus could replicate efficiently in mice, ferrets and dogs when the H6N1 AIV caused a French Turkey outbreak. At the same time a H6 antibody was detected in human serum [22, 40, 42-44]. In May 2013, the H6N1 virus was isolated from a 20 year old female in Taiwan. The sequence analysis of her H6N1 virus showed that the strain was highly homologous to the local avian influenza virus [23, 45]. In order to further research the outbreak risks and evolutionary patterns of the LPAIV H6N1 virus we conducted our research.

The Poyang Lake is located in Eastern China, yet it is not part of the West Asia-East Africa migration route. The results of the present study indicate that a new H6N1 isolate virus was produced via the recombination of different viruses in the natural gene pool. The figures 1-3 results demonstrated that gene fragments of the H6NI strain were derived from different migratory routes of birds. Therefore, there is a genetic interaction between different migration routes of birds in China. The data also showed, the H6N1 gene isolates were part of the Group III virus cluster of the natural gene pool, which evolved from the European library gene.

For our study we analyzed eight gene segments of the H6NI and reviewed other similar studies. The NA and M gene fragments we analyzed from the Republic of Georgia and Israel may have been transmitted by wild birds migrating in the West Asia-East Africa channel.

We also analyzed the NS1 gene and found it similar to the 2009 H6N5 gene in Korea and the A/domestic green-winged teal/Hunan/2036/2007(H3N6). Furthermore we found our NS1 gene similar to the 2011 H6N1 virus located in gray ducks of the National Nature Reserve in Jilin Province. According to the collected data the NS1 gene may exist as a “reserve gene” in the natural gene pool of the East Asia-Australia migration route. In addition the PB1 gene we analyzed was similar to 10 strains of the 2013 H10N5, H10N7 and H7N7 wild bird Jiangxi Province subtype samples. Moreover, we analyzed NP genes and found it similar to the 2013 H7N7 and H10N3 viruses in Jiangxi. Furthermore it was 99% similar to the 2011 H1N2 and 2012 H12N8 isolates from Dongting Lake. However since 2010 in China there hasn’t been any reported of similar gene sequences to the M, NA and HA genes. Since the H6N1 virus needs a period of integration in order to produce a new recombinant virus. Based on the discussed data and our NA evolution analysis it is revealed that H6N1 isolates may be produced by virus recombination in the migration process of Southeast Asia H6 subtype virus and wild birds from the Republic of Georgia.

For this study the H6N1 isolate gene and protein fragments were derived from the East Asia - Australia and West Asia - East African migratory flyway. It is speculated that the virus was brought into Western China by migratory birds from the West Asia-East Africa migratory route. When the infected birds stopped in gathering locations in Western China the H6N1 virus was transferred to other birds. Many of those birds then migrated towards Southern China and spread the H6N1 virus there. The migratory birds can converge and integrate virus genes from different sites thus promoting and accelerating new recombinant viruses. In conclusion new strains were created as a result of gene exchanges between different migratory routes. For example during our AIV monitoring we found the existence of LPAIV in the natural gene pool. The LPAIV has undergone continuous evolution and recombination in the natural gene pool of China. The continues development of the LPAIV can provide gene fragments for the HPAIV.

The phylogenetic analysis results revealed that the H6N1 isolates, the G1-like H9N2, the CK/Bei H9N2 and the Gs/GD-like H5N1 have different evolutionary patterns. However the A2837 virus infection in Taiwan has internal genes from the H5N1 virus, W312-like H6N1 and H6N1.

Our analyses of the isolated H6N1 virus strain revealed that it binds to the alpha-2, 3-sialic acid receptor of avian. Surprisingly we discovered that our isolated H6N1 virus could cause interstitial pneumonia in mice. Furthermore our site analysis revealed that the PB2, M and NS genes had mutations that increased the polymerase activity of the strain and enhanced the pathogenicity of the virus in mice. In conclusion we discovered that the LPAIV can be a potential threat to mammalian health and can provide gene fragments for the HPAIV. We can help prevent and control the threat of the H6NI virus by studying its evolution, pathogenic strains, and mutated genes that can endanger both humans and birds. Therefore by actively monitoring the AIV we can find its dynamic changes and provide the basis for further research on AIV’s origin and evolution.

## Acknowledgments and Funding

The project was supported by grants from National key R&D program of China, No. 2017YFD0501702, China Agriculture Research System Poultry-related Science and Technology Innovation Team of Peking (CARS-PSTP), the National Natural Science Foundation of China (No.81101835).

